# Structurally distinct duplex telomere repeat-binding proteins in *Ustilago maydis* execute specialized, non-overlapping functions in telomere recombination and telomere protection

**DOI:** 10.1101/2020.11.06.371997

**Authors:** Eun Young Yu, Syed Zahid, Min Hsu, Jeanette Sutherland, William K. Holloman, Neal F. Lue

**Affiliations:** Department of Microbiology & Immunology, W. R. Hearst Microbiology Research Center, Weill Medical College of Cornell University, New York, New York, United States of America; Sandra and Edward Meyer Cancer Center, Weill Cornell Medicine, New York, New York, United States of America

## Abstract

Duplex telomere binding proteins exhibit considerable structural and functional diversity in different phyla. Herein we address the distinct properties and functions of two Myb-containing, duplex telomere repeat-binding factors in *Ustilago maydis*, a basidiomycete fungus that is evolutionarily distant from the standard budding and fission yeasts. The two telomere-binding proteins in *U. maydis*, named *Um*Trf1 and *Um*Trf2, have different domain organizations and belong to distinct protein families with different phylogenetic distributions. Despite these differences, they exhibit comparable affinities and similar sequence specificity for the canonical, 6-base-pair telomere repeats. Deletion of *trf1* triggers preferential loss of long telomere tracts, suggesting a role for the encoded protein in promoting telomere replication. Trf1 loss also partially suppresses the ALT-like phenotypes of *ku70*-deficient mutants, suggesting a novel role for a telomere protein in stimulating ALT-related pathways. In keeping with these ideas, we found that purified Trf1 can modulate the helicase activity of Blm, a conserved telomere replication and recombination factor. In contrast, *trf2* appears to be essential and transcriptional repression of this gene leads to severe growth defects and profound telomere aberrations that encompass telomere length heterogeneity, accumulation of extrachromosomal telomere repeats such as C-circles, and high levels of single-stranded telomere DNA. These observations support a critical role for *Um*Trf2 in telomere protection. Together, our findings point to a unique, unprecedented division of labor between the two major duplex telomere repeat-binding factors in *Ustilago maydis*. Comparative analysis of *Um*Trf1 homologs in different phyla reveals a high degree of functional diversity for this protein family, and provides a case study for how a sequence-specific DNA binding protein can acquire and lose functions at different chromosomal locations.

---

Eukaryotic chromosome ends, or telomeres, harbor repetitive DNA sequences that mediate the assembly of a special, dynamic nucleoprotein structure that is crucial for genome stability. This dynamic structure stabilizes chromosome ends by suppressing aberrant repair events at the termini ^1, 2^, and by promoting the retention and replenishment of telomere DNA through successive rounds of replication. Special mechanisms are required to retain and replenish telomere DNA owing to (1) the propensity of telomere DNA to form barriers that block replication forks ^3^; and (2) the “end-replication” problem that prevents the complete synthesis of lagging strand duplex ^4^. Both the protective and DNA maintenance functions of telomeres are mediated by specific proteins within the dynamic assembly, often through direct physical interactions between telomere proteins and functionally relevant downstream targets.

Within the dynamic telomere nucleoprotein assembly, duplex telomere repeat binding factors are known to execute crucial functions ranging from telomere length regulation, telomere protection and telomere replication. Studies in diverse organisms have revealed considerable structural and functional variabilities in these duplex-binding factors. In mammals, the two major duplex telomere-binding proteins, TRF1 and TRF2, are each part of the shelterin complex that collectively protects chromosome ends and suppresses DNA damage response. While TRF1 and TRF2 are structurally similar in having an N-terminal TRFH dimerization domain and a C-terminal Myb domain, each protein serves distinct functions in telomere regulation. TRF2, in particular, plays a key role in telomere protection by inhibiting telomere fusions and the DNA damage response ^2^. TRF1, on the other hand, promotes efficient telomere replication by recruiting BLM helicase to unwind replication barriers ^5^. In the fission yeast *S. pombe*, a single TRF1/2-like protein named Taz1 alone performs multiple functions in telomere protection and replication, although it is unclear how the replication function is executed ^6, 7^. Interestingly, *S. pombe* contains two other Myb-containing proteins capable of binding telomere repeat-like sequences. One of these, named Tbf1, structurally resembles mammalian TRF1/2 (in fact, more so than does Taz1), but exhibits no compelling telomere functions ^8^. The other protein, named *Sp*Tay1/Teb1, bears two consecutive Myb motifs away from the C-terminus, and evidently functions primarily as a transcription factor ^9^(Fig. 1). Interestingly, the Tay1 homolog in the budding yeast *Yarrowia lipolytica* (the founding member of this protein family) is reported to be the main telomere-binding protein and plays a critical role in telomere maintenance and protection ^10^. While the evolutionary basis for the structural and functional divergence of duplex telomere binding proteins is not fully understood, a likely driver is the irregularity of the fission and budding yeast telomere repeats (e.g., TTAC _0-1_A _0-1_C _0-1_G_1-9_ for *S. pombe* and TG_1-3_ for *S. cerevisiae*) and its deviation from the canonical GGGTTA repeat found in mammals and all major eukaryotic branches. For a more detailed discussion of telomere sequence divergence in budding and fission yeasts and the corresponding changes in duplex telomere binding proteins, see Sepsiova et al. and Steinberg-Neifach and Lue ^11, 12^.

**Figure 1.**
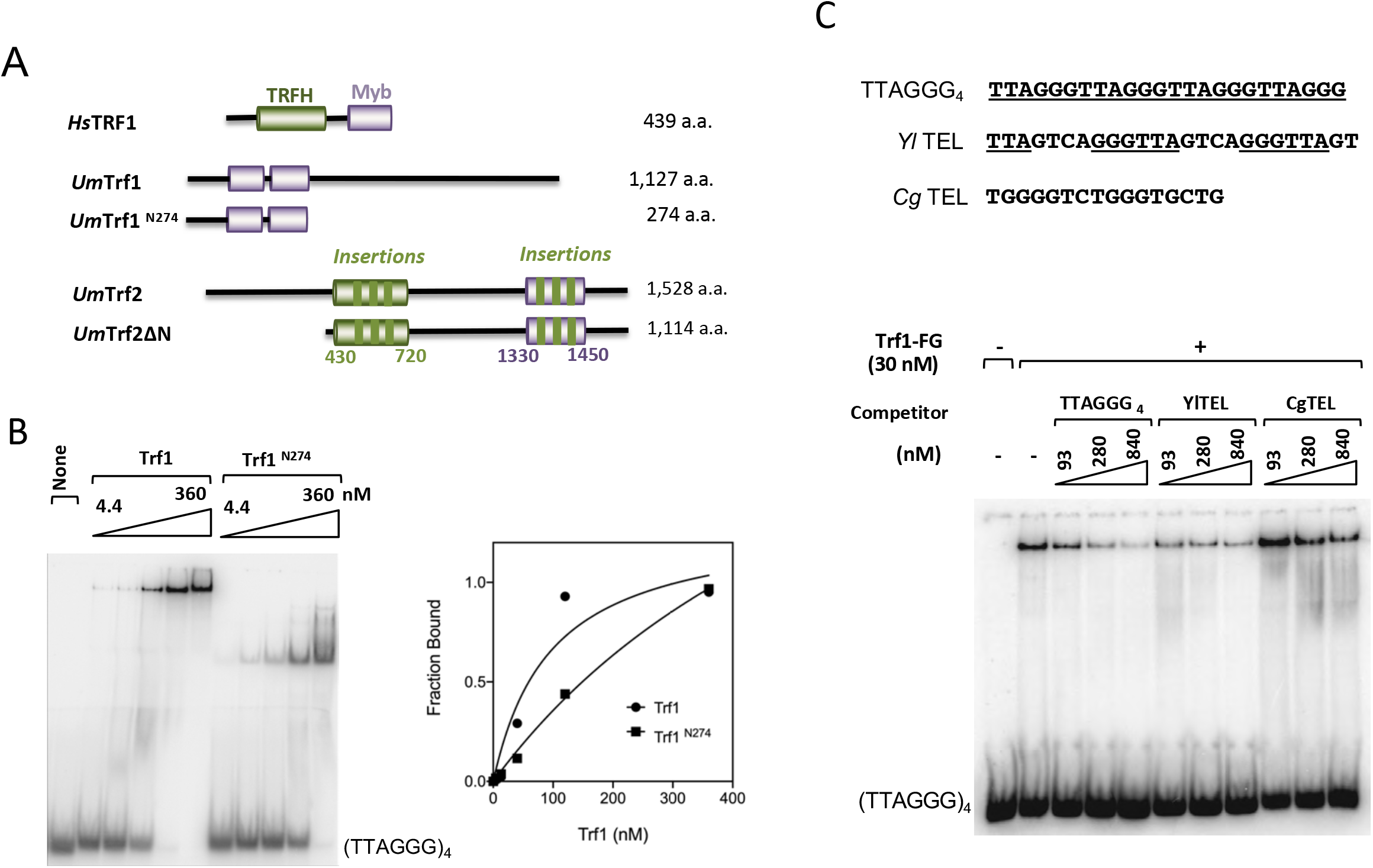
The DNA-binding activity of *Ustilago maydis* Trf1. (A) Schematic illustrations of the domain structures of the Ustilago maydis Trf1 and Trf2 proteins analyzed in this study, as well as that of human TRF1, a prototypical member of the TRF/TBF1 protein family. The TRFH and Myb domains are shown in green and purple, respectively. (B) EMSA analysis of the DNA-binding activities of Trf1 and Trf1^N274^. A representative series of assays is shown on the left and the quantitation shown on the right. (C) The DNA-binding specificity of Trf1 was examined using three different double-stranded competitor oligonucleotides. The sequences of the competitors are shown on top with the canonical GGGTAA repeat unit underlined – only the G-strand sequences are displayed. A representative set of assays is shown on the bottom.

A potentially informative model with which to address mechanistic and evolutionary issues related to duplex telomere binding proteins is the basidiomycete *Ustilago maydis*. This model was originally developed by Robin Holliday to investigate recombinational repair ^13^. We were initially attracted to this model – in comparison to other fungal models – because (1) *U. maydis* carries a telomere repeat unit that is identical to the mammalian repeat ^14^; (2) the recombination and repair machinery in this organism bears greater resemblances to those in mammals ^15^; and (3) repair proteins in this organism (e.g., Rad51, Blm and Brh2) promote telomere replication and telomere recombination just like their mammalian ortholog counterparts ^16, 17, 18^. Accordingly, we have taken advantage of *U. maydis* to examine the mechanisms and regulation of repair proteins at telomeres ^16, 17, 18, 19^. Interestingly, with respect to duplex telomere binding proteins, our initial database query (as well as a previous bioinformatics screen) revealed in the *U. maydis* genome just one potential duplex telomere binding protein that is structurally similar to *Sp*Tay1 and that we and others named *Um*Trf1 ^11, 19, 20^. We confirmed that this protein has high affinity and sequence specificity for the cognate telomere repeats, and that it interacts directly with Blm helicase, just like mammalian TRF1/2 ^19^. While these initial findings imply that *Um*Trf1 could be the main duplex telomere repeat factor in *U. maydis*, subsequent analysis of *trf1Δ* mutants revealed little telomere de-protection phenotypes (see below), suggesting that the true ortholog of mammalian TRF2 remained to be identified. Indeed, Tomaska and colleagues have noted the existence of a TRF/TBF1-like gene *in U. maydis* with unknown function and potential TRFH and Myb domains (Uniprot ID: A0A0D1E1Z3)^11^. Thus, *U. maydis* might harbor two structurally distinct, Myb-containing proteins that mediate distinct or overlapping functions in telomere protection and maintenance. Because basidiomycetes occupy a more basal position in fungal phylogeny than the ascomycete budding and fission yeasts, studies of duplex telomere-binding proteins in this lineage may offer insights on the origin of these protein families in fungi and the division of labor between them vis-à-vis other organisms.

Herein we report a structural and functional analysis of the two duplex telomere repeat binding proteins in *U. maydis*, which we named Trf1 (Tay1-like) and Trf2 (TRF and TBF1-like). We showed that both proteins, despite their unusual structural features (see below), recognize the canonical telomere repeat GGGTTA/TAACCC with high affinity and sequence specificity. Trf2 but not Trf1 is critical for telomere protection. Trf2 appears to be essential and transcriptional repression of this gene results in profound telomere aberrations, accumulation of ssDNA, and elevated levels of extrachromosomal telomere repeats. Trf1, on the other hand, contributes to telomere replication and positively stimulates an ALT-like pathway of telomere recombination, most likely by interacting with Blm and controlling its helicase activity. This functional division of labor is unlike that observed in metazoans or in budding/fission yeasts. Cross comparison of our results with those from other fungal and non-fungal systems highlights the potential of structurally distinct telomere proteins to execute similar functions in different lineages. Based on these results and others, we propose that the emergence of the Tay1 family of telomere proteins in fungi may have provided added flexibility to the fungal telomere system to adapt to drastic changes in telomere sequences in budding and fission yeasts.

## Results

### *U. maydis* Trf1 binds GGGTTA/TAACCC with high affinity and sequence specificity through its N-terminal, duplicated Myb motifs

*Um*Trf1 is distinguished structurally from both the *Y. lipolytica* and *S. pombe* Tay1 homologs in having a much longer C-terminal extension (following the tandem Myb motifs), which renders the protein twice as large as the other homologs. To investigate the potential function of the *Um*Trf1 C-terminus in DNA-binding, we purified an N-terminal 274 amino acid (a.a.) fragment (spanning the duplicated Myb motifs and named TRF1^N274^) and compared its DNA-binding properties to those of the full length protein (Fig. 1A). In a titration analysis, TRF1 exhibited a slightly higher affinity binding to telomere DNA than Trf1^N274^; the estimated K_d_s of these proteins for the TTAGGG_4_ probe are 35 and 110 nM, respectively (Fig. 1B). We also assessed the sequence-specificity of Trf1 using competitor oligonucleotides harboring different telomere repeats. The *Yl*TEL oligo, in which the canonical 6-bp repeat unit (GGGTTA) is interspersed with other sequence, competed as effectively as the cognate 6-bp repeat (TTAGGG_4_) (Fig. 1C). In contrast, the *Cg*TEL oligo, another G-rich repeat that lacks the GGGTTA sequence, was unable to compete. This binding preference is similar to that reported for *Y. lipolytica* Tay1, specifically with regard to the strong preference for DNA with canonical repeats ^10, 11^, and indicates that *Um*Trf1 has the requisite DNA-binding properties to act specifically at *U. maydis* telomeres.

### *U. maydis Trf2* is also a high affinity telomere repeat binding factor

In addition to *Um*Trf1, Sepsiova et al. noted the potential existence of a TRF/TBF1-like protein (Uniprot ID: A0A0D1E1Z3), which, being 1,528 a.a. long, is considerably larger than a typical TRF/TBF1 ortholog such as human TRF1 (Fig. 2A). We named this gene *Umtrf2*, and performed additional bioinformatic analysis by identifying and characterizing closely related homologs in the Pucciniomycotina and Ustilaginomycotina subphyla (Supp. Fig. 1A and 1B). Like *Um*Trf2, these fungal proteins are ^~^1.5 – 3 times the sizes of mammalian TRFs. Nevertheless, multiple sequence alignment of these proteins against metazoan TRF orthologs revealed the existence of both a central TRFH domain and a C-terminal Myb domain, precisely as predicted for a true TRF ortholog (Supp. Fig. 1A and 1B). Notably, the fungal proteins carry multiple insertions in both their TRFH and Myb domains (vis-à-vis the metazoan proteins), thus accounting in part for their larger sizes. Notwithstanding these multiple insertions, structural considerations suggest that the fungal protein domains may retain the known function of TRFH and Myb in dimerization and DNA-binding. For example, even though the Myb motif of *Um*Trf2 harbors three short insertions (Fig 1A and Supp. Fig. 1B), these insertions map to regions of the motif that are distal to the DNA-binding surface as judged by sequence alignment and structure of the human TRF1-DNA complex (Fig. 2A). Accordingly, they are not predicted to disrupt *Um*Trf2-DNA binding. Notably, both insertion 2 and 3 are relatively invariant in length and contain well conserved residues, suggesting that they may serve significant functions that are unrelated to DNA-binding.

**Figure 2.**
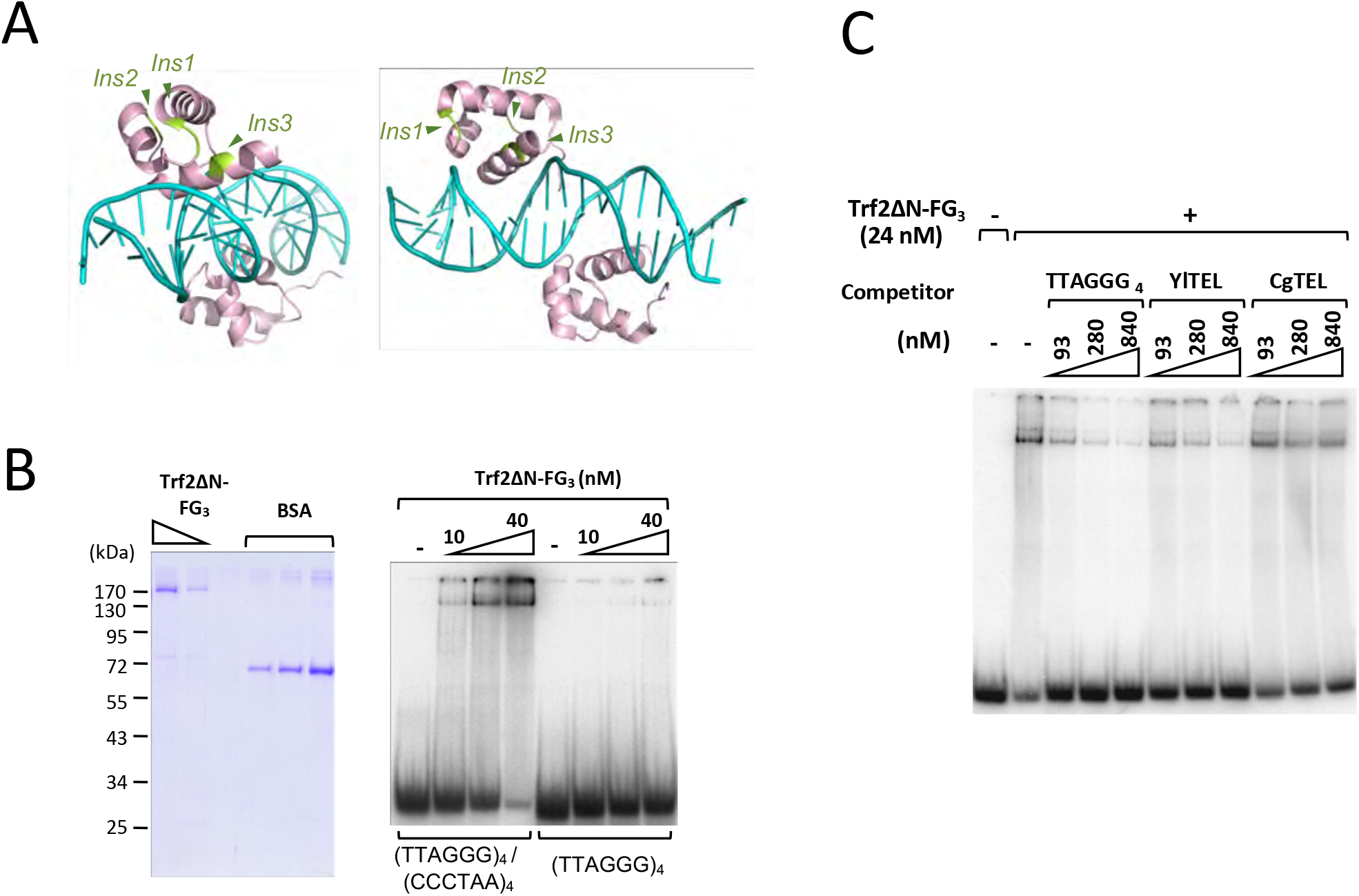
The DNA binding activity of *Ustilago maydis* Trf2. (A) The structure of human TRF1-DNA complex is used to illustrate the locations of three insertions within the Myb domain of *Ustilago maydis* Trf2. The positions where the insertions occur are shown in light green. Note that these are all positioned away from the DNA-protein interface. (B) (Left) Purified Trf2ΔN-FG_3_ was analyzed by SDS-PAGE along with BSA standards. (Right) EMSA analysis was performed using varying concentrations of Trf2ΔN-FG_3_ and either single-stranded or double-stranded telomere oligonucleotides. (C) The DNA-binding specificity of Trf2 was examined using the indicated competitor oligonucleotides.

To test the DNA-binding activity of *Um*Trf2 directly, we purified and analyzed an N-terminally truncated form of the protein (the full length protein did not express well in *E. coli*), and showed that this truncated protein (Trf2ΔN-FG_3_) binds with high affinity to double stranded TTAGGG_4_ (K_d_ = 20 nM) (Fig. 2B). Moreover, competition experiments indicate that just like *Um*Trf1, *Um*Trf2 strongly prefers DNAs that contain the canonical 6-bp repeat (Fig. 2C). Further analysis indicates that the two putative telomere-binding proteins have nearly indistinguishable minimal DNA target sizes of ^~^ 2—3 repeats (Supp. Fig. 2A and 2B). A probe consisting of 2.5 repeats was bound with lower affinity by both Trf1 and Trf2, and a probe with just 2 repeats did not support any detectable binding. Thus, despite the structural differences both within and outside the Myb domain of Trf1 and Trf2, the two *U. maydis* proteins evidently recognize telomere repeat DNA using very similar mechanisms.

### U. maydis Trf1 is required for optimal telomere maintenance, but not telomere protection

To investigate the telomeric functions of Trf1 *in vivo*, we analyzed the phenotypes of multiple, independently derived *trf1Δ* clones. These clones were found to exhibit similar growth rates as the parental strain (FB1), but no evidence of senescence following prolonged passages, suggesting that Trf1 is not essential for either telomere protection or telomerase function (data not shown). However, telomere restriction fragment (TRF) Southern analysis revealed substantially shortened but stably maintained telomeres in the *trf1Δ* mutants (Fig. 3A). The shortening was evident at streak 3 (corresponding to ^~^100 generation after the construction of the mutants), but did not worsen at later time points. Indeed, both wild type and *trf1Δ* telomeres grew slightly during continuous sub-culturing, a phenomenon that has been noted earlier ^17^. The telomere phenotypes of *trf1Δ* clones are reminiscent of DNA repair mutants and point to telomere replication as a potential function of Trf1 ^17, 19^. Previous characterization of the telomere replication mutants (e.g., *blmΔ* and *rad51Δ*) revealed preferential loss of long telomeres (^21^ and data not shown). This feature is also true of *trf1Δ* telomeres: in STELA analysis of UT4-bearing telomeres, the longest telomere tracts found in *trf1Δ* were ^~^300 bp shorter than those in the wild type strain, whereas the shortest tracts were similar in size (Fig. 3B). (UT4 is a previously characterized subtelomeric element in *U. maydis* ^14, 21^.) While the telomere phenotype of *trf1Δ* resembles that in the DNA repair mutants, we found that *trf1Δ* did not manifest significantly greater susceptibility to various DNA-damaging agents than the parental strain, except for a slight sensitivity to MMS (Fig. 3C, bottom panel).

**Figure 3.**
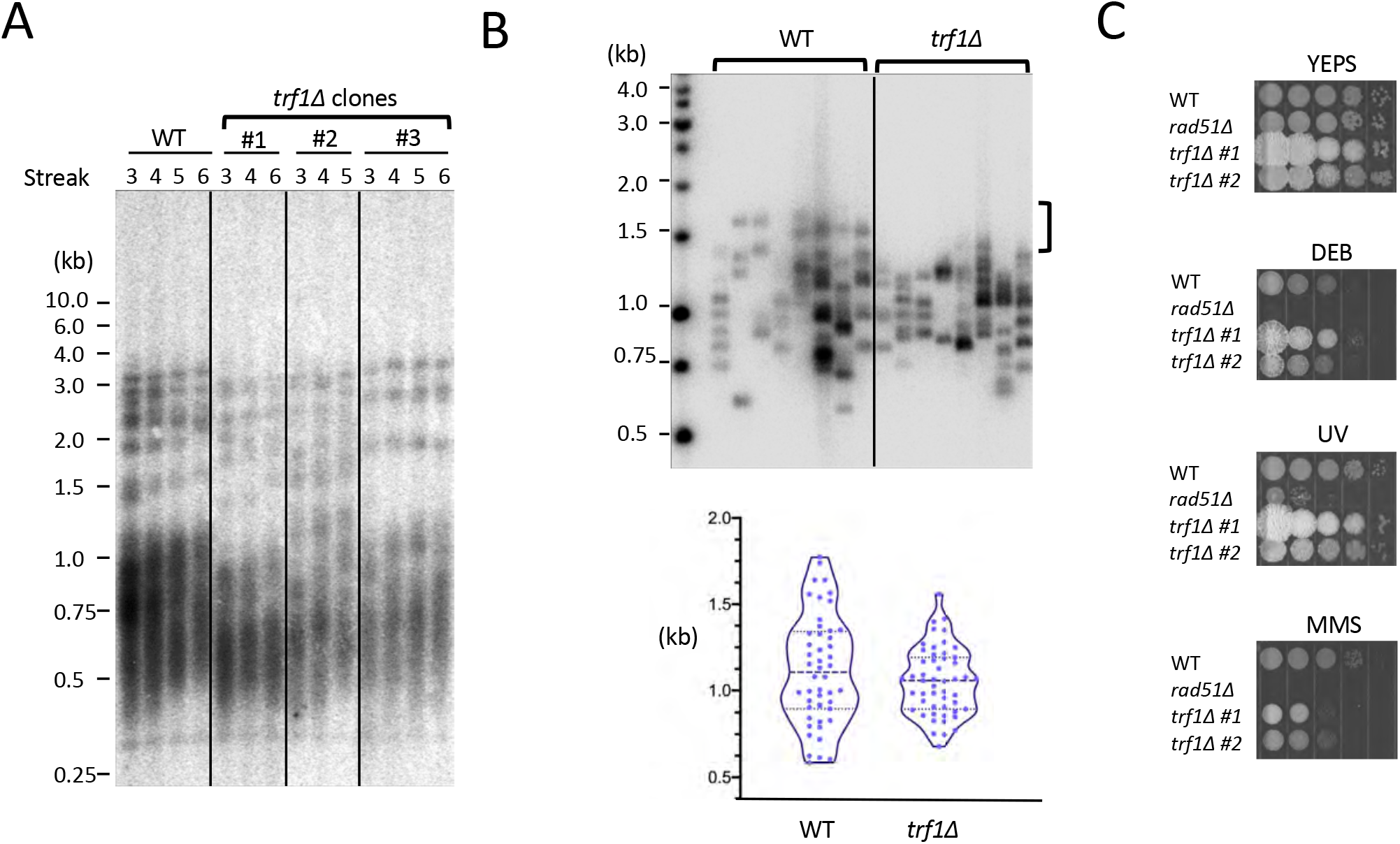
*Ustilago maydis* Trf1 is required for optimal telomere replication, and plays a minor role in DNA damage response. (A) Chromosomal DNAs were prepared from wild type and three independent *trf1Δ* transformants following the indicated number of streaks. Following *Pst*I cleavage, the DNAs were subjected to TRF Southern analysis. (B) Chromosomal DNAs from wild type and *trf1Δ* strains were subjected to STELA. Following ligation to the Teltail oligonucleotide, UT4/5-containing telomeres were amplified using a UT4/5-specific primer, and detected using a UT4/5 probe. Representative assays are shown on the top, with the vertical bracket to the right calling attention to the long telomere fragments missing from the *trf1Δ* samples. The composite STELA profiles are displayed as violin plots on the bottom. (C) Ten-fold serial dilutions of the indicated strains were spotted on YPD containing the indicated clastogens (0.02% MMS Methyl methanesulfonate; 0.01% DEB, 1,2,3,4-diepoxybutane) or subjected to UV irradiation (120 J/m^2^), and their growth assessed after 2-3 days.

In contrast to the telomere shortening observed in Southern and STELA analysis, in-gel hybridization assays did not uncover significant changes in ssDNA at *trf1Δ* telomeres (data not shown). In addition, we did not detect any C-circles, which are extrachromosomal circular telomere repeats that often accumulate when deprotected telomeres become recombinogenic (data not shown) ^18^, ^22^. Thus, the predominant telomere function of *Um*Trf1 seems to be limited to telomere replication.

### *U. maydis* Trf1 interacts directly with Blm helicase and modulates its helicase activity

We have previously shown that just like mammalian TRF1 and TRF2, *Um*Trf1 physically interacts with the cognate Blm helicase ^19^. Given this interaction and the similarity between the telomere phenotypes of *trf1Δ* and *blmΔ*, we hypothesize that Trf1 promotes telomere replication by controlling Blm activity. To test this possibility biochemically, we analyzed the effects of Trf1 on Blm helicase activity *in vitro* (Fig. 4A). Two double-stranded DNA substrates with 3’ overhangs, one carrying a telomeric sequence, and the other a non-telomeric sequence, were used to assess the helicase activity of Blm, a 3’ to 5’ helicase ^23^. Notably, the untreated telomeric substrate migrated as two distinct species in native gels, most likely owing to G-rich overhang-mediated dimer formation (Fig. 4B, leftmost lane; Supp. Fig. 3, 6^th^ lane). As expected, the addition of Blm resulted in the unwinding of duplex DNA into ssDNA in an ATP-dependent manner (Supp. Fig. 3). Interestingly, the effects of Trf1 on unwinding were substrate sequence-dependent: on telomeric DNA, Trf1 inhibited DNA unwinding by Blm by up to 70%, whereas on non-telomeric DNA, the effect was stimulatory by up to 2.5 fold (Fig. 4A and 4B). The inhibitory effect of full length Trf1 can be recapitulated by the N-terminal Myb domain (Trf1 ^N274^), suggesting that high affinity DNA-binding by Trf1 alone may be sufficient to block the unwinding activity of Blm (Supp. Fig 4B). In contrast, Trf1 ^N274^ is unable to stimulate unwinding on the non-telomeric substrate, implying that the C-terminal regions of Trf1 contribute to this activity, possibly through protein-protein interactions (Supp. Fig. 4A).

**Figure 4.**
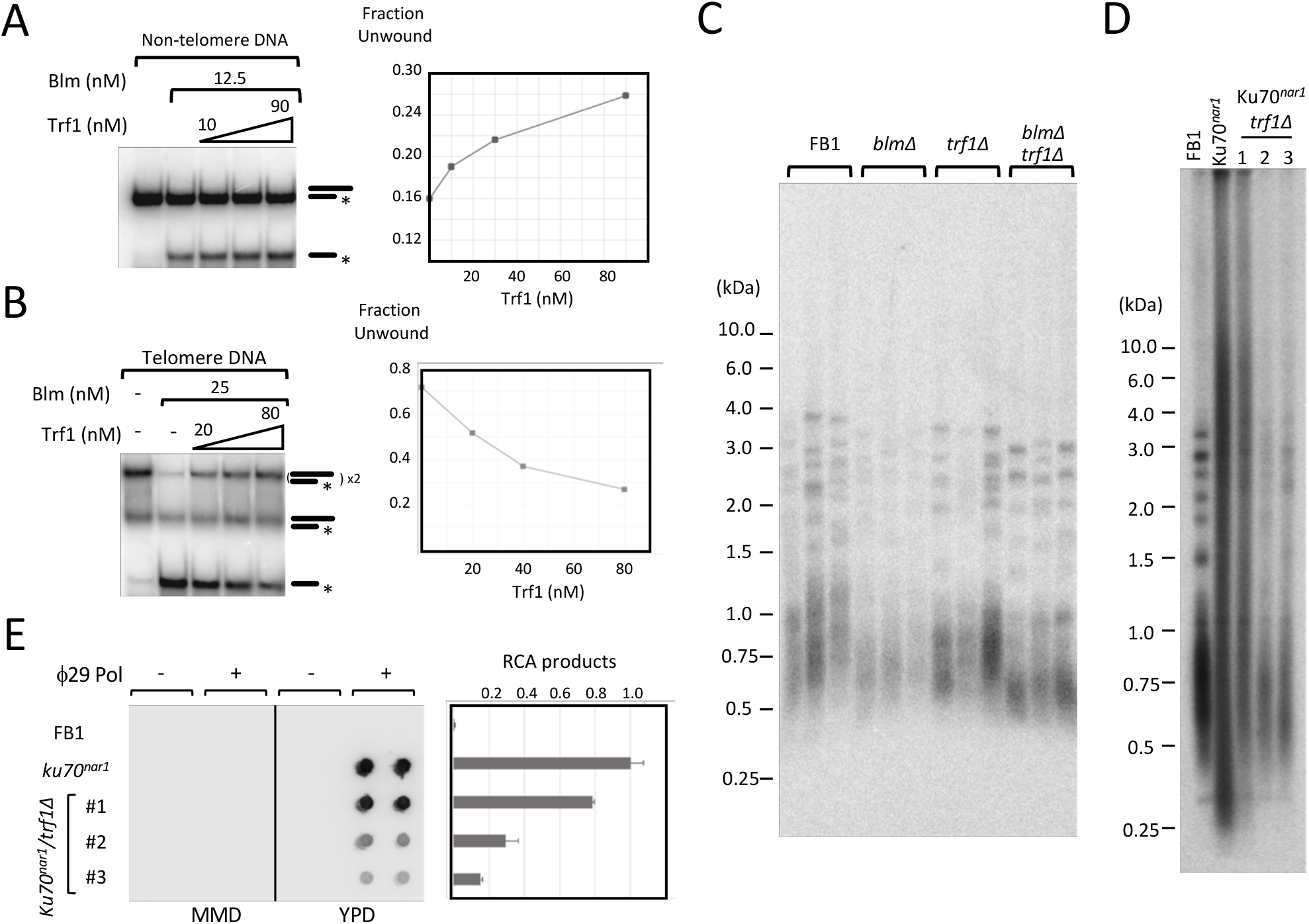
Trf1 modulates Blm helicase activity and functionally resembles Blm in regulating telomere replication and recombination. (A) The helicase activity of Blm (12.5 nM) on a non-telomeric substrate was examined in the presence of increasing Trf1 concentrations (0, 10, 30 and 90 nM). The quantified results are shown on the right. (B) The helicase activity of Blm (25 nM) on a telomeric DNA-containing substrate was examined in the presence of increasing Trf1 concentrations (0, 20, 40 and 80 nM). The quantified results are shown on the right. (C) Chromosomal DNAs were isolated from three independent cultures of wild type or mutant strains and subjected to telomere restriction fragment analysis. The mutant strains were all passaged for ^~^ 75 generations before the analysis. (D) Chromosomal DNAs were isolated from the indicated strains (grown in YPD), and subjected to telomere restriction fragment Southern. (E) Chromosomal DNAs were isolated from the indicated strains with the given growth condition, and subjected to duplicate C-circle assays. The scan of the blot is shown on the left and the quantification (normalized against the average value of *ku70^nar1^* samples) plotted on the right.

To examine functional interactions between *Um*Trf1 and Blm *in vivo*, we generated *blmΔ* and *trf1Δ blmΔ* mutants in the same strain background as *trf1Δ* and compared their telomere phenotypes. All three mutants manifested stably shortened telomeres, consistent with a telomere replication defect (Fig. 4C). Notably, as judged by the loss of long telomeres in both telomere Southern and STELA analysis, the defect is more severe in *trf1Δ blmΔ* than in either single mutant (Fig. 4C, Supp. Fig. 5). These results suggest that the two genes do not act exclusively in a single pathway in promoting telomere replication. Indeed, we had previously detected an interaction between *U. maydis* Blm and Pot1, raising the possibility that Blm may be regulated by multiple telomere proteins ^19^. Likewise, it is possible that Trf1 may interact with another factor(s) that contributes to telomere replication. However, with respect to potential interacting partners, we note that Trf1 does not appear to bind Rad51/Brh2, another DNA repair complex implicated in telomere replication ^17^(Suppl. Fig. 6).

Another previously established function of Blm at telomeres is to trigger an ALT-like pathway when telomeres are de-protected. More specifically, when the *ku70/80* complex is transcriptionally repressed in *U. maydis* (e.g. in the *ku70^nar1^* mutant), telomeres become highly aberrant and manifest a collection of abnormalities that resemble those in ALT cancer cells, including telomere length heterogeneity, ssDNA on the C-strand, and high levels of C-circles ^16, 18^. All these phenotypes are completely suppressed in *ku70^nar1^ blmΔ*, implying an essential role for *blm* in triggering the ALT pathway. Given the ability of Trf1 to regulate Blm activity, we proceeded to generate *ku70^nar1^ trf1Δ* mutants and analyzed their telomeres. Notably, all three independently generated *ku70^nar1^ trf1Δ* exhibited milder telomere length abnormalities than *ku70^nar1^* when *ku70* expression was repressed. In particular telomere length distributions became less heterogeneous, and the levels of C-circles were substantially reduced in *ku70^nar1^ trf1Δ* relative to that in *ku70^nar1^* (Fig. 4D and 4E). The levels of C-strand ssDNA in the double mutants were also significantly lower than that in the *ku70^nar1^* single mutant (Supp. Fig. 7A). In accordance with the mild telomere aberrations, the *ku70^nar1^ trf1Δ* double mutants exhibited very little growth defects and did not manifest the enlarged and elongated morphology characteristic of G2/M arrest (Supp. Fig. 7B and 7C). These results indicate that *trf1Δ* partially phenocopies *blmΔ* in the context of Ku70 depletion, and support the notion that Trf1 positively regulates Blm in activating the ALT pathway. The residual telomere aberrations in the *ku70^nar1^ trf1Δ* mutants show, however, that Trf1 is not absolutely essential for Blm function in this pathway.

### *U. maydis trf2* is crucial for telomere protection

The lack of an obvious telomere deprotection phenotype in *trf1Δ* mutants suggests that *Um*Trf2 may be the key telomere protection factor. To test this idea, we attempted to generate null mutants by gene replacement. Multiple attempts were unsuccessful, suggesting that the gene may be essential (data not shown). We then constructed transcriptionally conditional mutants of *trf2* (named *trf2^crg1^*) in which the endogenous *trf2* promoter is replaced by that of the regulable, arabinose-dependent promoter *crg1* ^24^. As a consequence, *trf2* expression in these mutants is induced in YPA but repressed in YPD. Notably, all the mutant clones grew poorly in YPD (albeit with some differences in growth rate), indicating that the protein is required for optimal cell proliferation (Fig. 5A and data not shown). The mutants also manifest elongated morphology that is consistent with cell cycle defects (Fig. 5B). TRF Southern analysis revealed in the *trf2*-repressed clones grossly abnormal telomeres characterized by (1) high overall telomere repeat content, (2) telomere size heterogeneity, and (3) high levels of extremely short telomere fragments (Fig. 5C, marked by a vertical bracket). Notably, a large fraction of telomere DNA in the mutant probably exists as extra-chromosomal telomere repeats (ECTR), given that short telomere fragments were detected even in the absence of restriction enzyme digestion (Fig. 5D). All three aberrant telomere features noted above were evident in multiple, independently-generated *trf2^crg1^* clones, and during serial passages in YPD, implying that Trf2 is critical for telomere protection (Fig. 5E).

**Figure 5.**
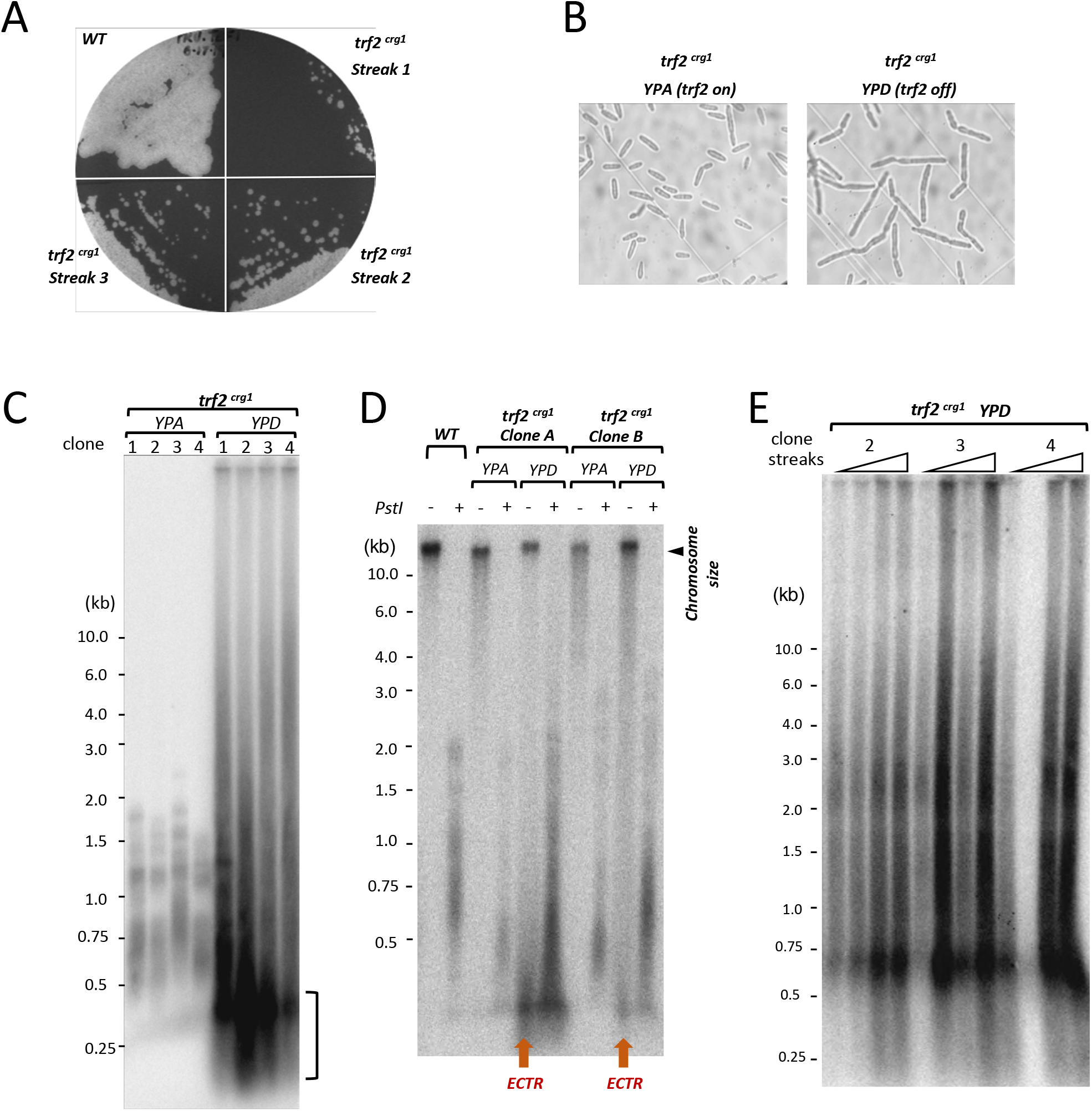
*Ustilago maydis* Trf2 is critical for cell proliferation and telomere protection. (A) A *trf2^crg1^* clone was passaged on YPD (a medium that represses *trf2* transcription), and the growth of the clone at successive streaks assessed along with the parental strain. The plates were incubated at 30°C for 3 days. (B) A *trf2^crg1^* clone was grown in liquid YPA or YPD, and examined under the microscope. (C) DNAs from multiple, independently constructed *trf2^crg1^* mutants that had been grown in YPA or YPD were subjected to TRF Southern analysis. (D) Untreated or *Pst*I-digested chromosomal DNAs were fractionated on agarose gel, transferred to nylon membrane, and then subjected to Southern hybridization using a (TTAGGG)x82 probe. The short telomere fragments that can be detected in the absence of *Pst*I digestion are highlighted as extra-chromosomal telomere repeats (ECTR). (E) Multiple independent *trf2^crg1^* clones were passaged on YPD plates by repeated re-streaking. DNAs from liquid culture out-growth of the resulting colonies were subjected to telomere restriction fragment Southern analysis.

The role of Trf2 in telomere protection is also supported by analysis of single-stranded telomere DNA through ingel hybridization. In particular, low levels of G-strand ssDNA (single-strand DNA) were detected in *trf2*-expressing clones, consistent with normal telomeres harboring G-strand, 3’-overhangs (Fig. 6A). In contrast, in *trf2*-repressed clones, G-strand ssDNA was significantly reduced whereas C-strand ssDNA became greatly elevated (Fig. 6A). Moreover, the amounts of C-circles, another marker of abnormal telomere recombination, were also increased in these clones (Fig. 6B). Notably, these ssDNA and C-circle aberrations are similar to those in *ku70* or *ku80*-repressed mutants, suggesting that the abnormal telomere processing reactions in the *trf2* and *ku*-deficient mutants may share some mechanistic resemblance. However, there are likely important differences as well, given that the levels of C-circles in the *ku70^nar1^* are more than 10-fold higher than those in *trf2^crg1^* (data not shown).

**Figure 6.**
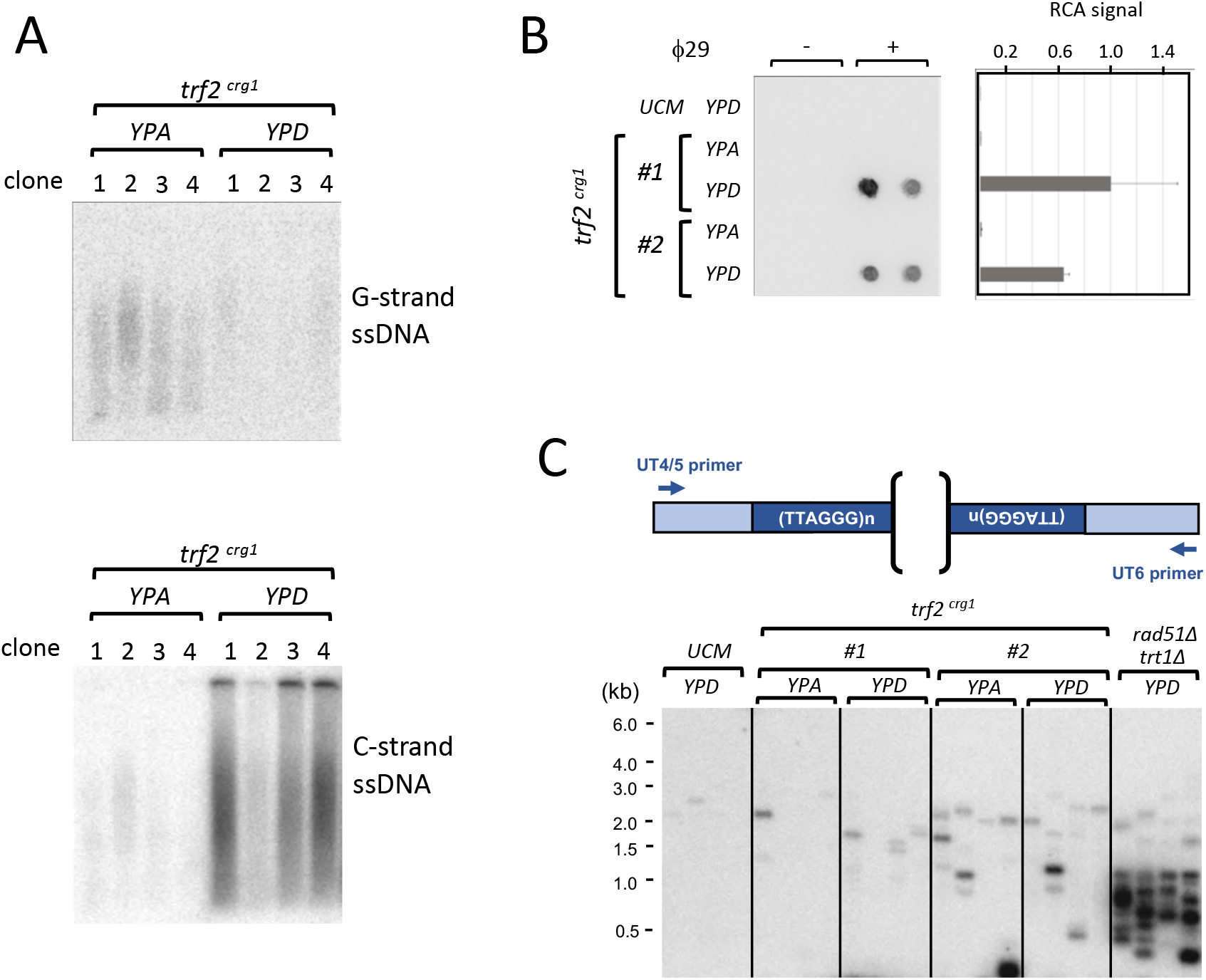
*Ustilago maydis* Trf2 suppresses multiple telomere aberrations. (A) DNAs from multiple, independently constructed *trf2^crg1^* strains grown in YPA or YPD were assayed for the levels of unpaired G-strand and C-strand telomere DNA by in-gel hybridization. (B) The indicated chromosomal DNA preparations were subjected to duplicate C-circle assays. The scan of the blot is shown on the left and the quantification (normalized to the average of the *trf2^crg1^* clone #1 YPD samples) plotted on the right. (C) The DNAs from the indicated strains grown in the given media were subjected to telomere fusion analysis.

**Figure 7.**
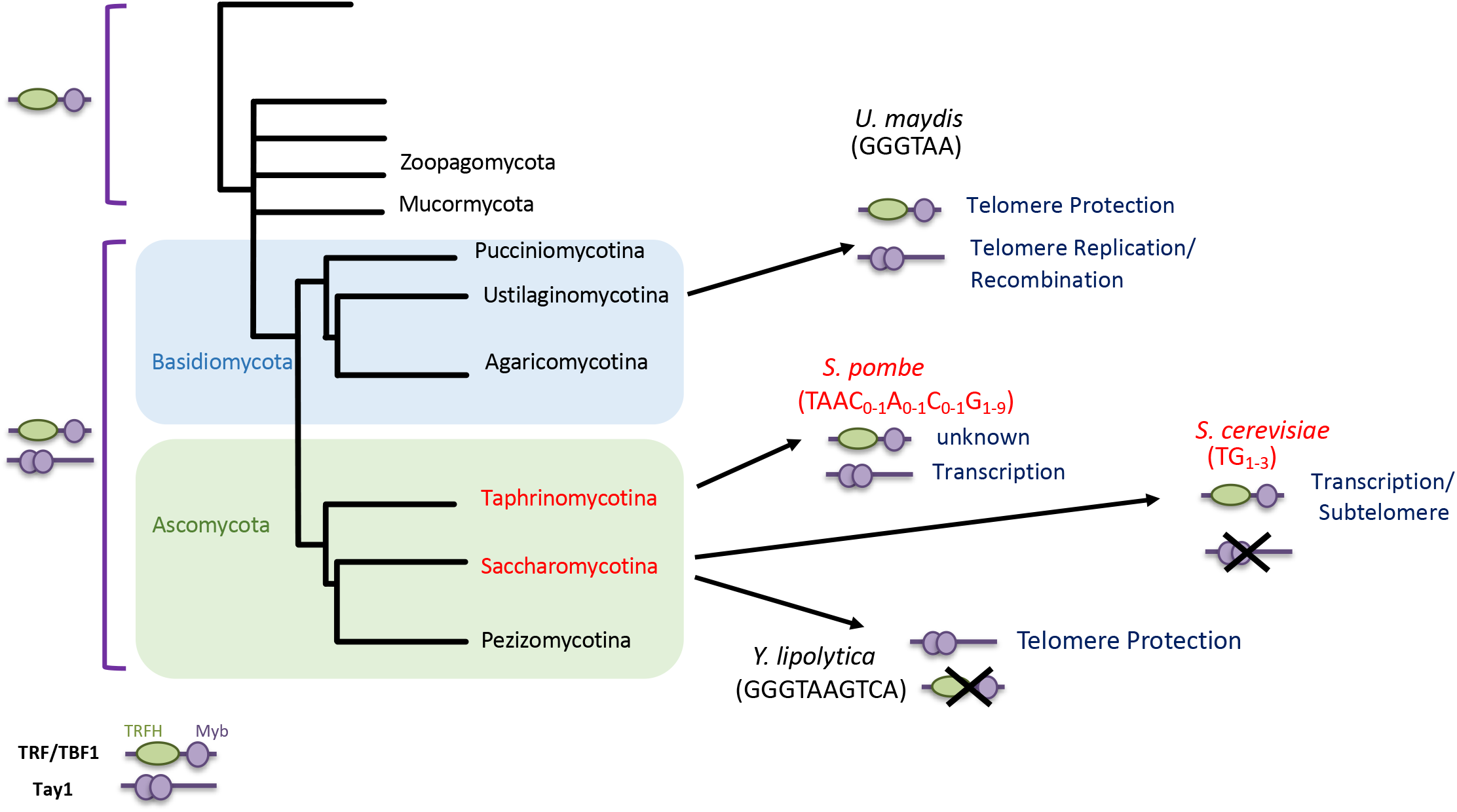
Functional plasticity of the Tay1 protein family. (Left) A simplified fungal phylogeny is shown along with the distribution of the TRF/TBF1 and Tay1 protein family in the phylogeny. The tree is adapted from a previous study ^48^. The phyla in which the telomere repeats deviate from the canonical GGGTAA sequence are indicated in red. (Right) The existence and functions of TRF/TBF1 and Tay1 homologs in selected organisms are illustrated.

To further examine the protective function of *trf2*, we developed a PCR-based assay to assess telomere-telomere (T-T) fusions between UT4/5 and UT6-bearing telomeres (Fig. 6B). Application of this assay to the *U. maydis trt1Δ rad51Δ* double mutant revealed high levels of T-T fusions, which are consistent with previous studies of such mutants in other organisms ^25, 26^. In contrast, the levels of fusions in the *trf2*-repressed clones are only slightly elevated, suggesting that the defective telomeres in these cells are not as prone to non-homologous end joining.

## Discussion

Fungi have been a rich source of discovery for telomere biology. Nevertheless, except for a few standard and atypical organisms in the ascomycete phylum, the telomere machinery in different fungal lineages remains poorly understood. We have sought to characterize telomere regulation in *Ustilago maydis* as a starting basis for understanding telomeres in basidiomycetes. Because of the resemblance of the repair machinery in *U. maydis* to that in metazoans, this fungus could also serve as a useful model for the interplay between the telomere and repair machinery. In this study, we focused on the structural, functional, and mechanistic features of two doublestrand telomere binding proteins in *U. maydis*, named *Um*Trf1 and *Um*Trf2. Most notably, we showed that both proteins, despite their unusual and quite different structural features, recognize the canonical telomere repeat GGGTTA/TAACCC with very similar affinity and sequence specificity. Remarkably, the two proteins mediate quite distinct telomere regulatory functions, with Trf1 contributing primarily to telomere replication and telomere recombination, and Trf2 being the chief telomere protection factor. These observations have interesting mechanistic and evolutionary ramifications, as discussed below.

### The DNA-binding mechanisms of the TRF/TBF1 and Tay1 protein families

The DNA-binding mechanisms of TRF/TBF1 protein family have been studied extensively. High resolution structures of the Myb domain-DNA complexes revealed crucial contacts between the two Myb protomers (head-to-tail arrangement) and consecutive GGG/CCC triplets in the telomere DNA targets ^27^. In addition, single particle EM analysis of full length TRF1 suggests that the TRFH dimerization domain helps to position the Myb domains in a relative orientation that is conducive to DNA binding ^28^. In contrast, small angle x-ray scattering (SAXS) envelope of TRF2 dimer revealed an extended conformation which leaves the two Myb protomers far apart ^29^. It seems likely that, given its similar structural organization and similar DNA-binding specificity, *Um*Trf2 would recognize telomere repeats in the same fashion as mammalian TRF1 and TRF2, i.e., through specific interactions between Myb protomers and two neighboring GGG/CCC triplets. One notable feature of *Um*Trf2 is the large linker between its TRFH and Myb domains (^~^600 amino acids compared to ^~^100 a.a. in human TRF1 and ^~^200 a.a. in human TRF2), a feature that could provide additional flexibility to *Um*Trf2 in positioning its two Myb protomers relative to the DNA target. This idea is consistent with the ability of *Um*Trf2 to bind *U. maydis* and *Y. lipolytica* telomere sequences with comparable affinities, despite the differences in gaps between the consecutive GGG/CCC triplets (3 bp for the *U. maydis* repeat and 7 bp for the *Y. lipolytica* repeat).

Less is known about the DNA-binding mechanism of the Tay1 protein family, which are characterized by two tandem Myb motifs separated by ^~^20 a.a (or much longer in some homologs such as those from *Coccidioides posadasii* or *Uncinocarpus reesei*) ^10, 19^. While no high resolution structural information is available for this protein family, the similarities in the DNA target size and sequence preference exhibited by *Um*Trf1 and *Um*Trf2 suggest that Tay1-like proteins may position their two Myb motifs on telomere DNA in the same way as TRF/TBF1 proteins. Notably, structural considerations suggest that the linker between the two Myb motifs in Tay1 (minimally 20 a.a.) is adequate to span the distance between the C-terminus of one Myb protomer and the N-terminus of the other protomer as judged by the crystal structure of the TRF1-DNA complex (30—40 Å depending on the combination). The long linker between the Myb motifs may also account for the ability of *Um*Trf1 to bind *Y. lipolytica* telomeres, in which the GGGTAA core repeat is separated from each other by 4 nucleotides. Even though the two Myb motifs are expected to be further apart along the DNA axis, they would lie on the same side of the DNA double helix, thus reducing the spatial separation that needs to be spanned by the protein linker. Therefore, we propose that a Tay1 monomer might be sufficient for high affinity and sequence-specific binding to telomere repeats. Indeed, our preliminary co-expression/pull down experiments did not reveal significant interactions between differently tagged *Um*Trf1 (Supp. Fig. 8), suggesting that the protein does not form a stable dimer. However, we cannot exclude the possibility of DNA-induced oligomerization, which was suggested by EM analysis of *Y. lipolytica* Tay1-DNA complex ^10^.

### The evolutionary plasticity of Tay1 function

The Tay1 protein family, which encompasses *Um*Trf1, provides an interesting case study in the plasticity of the telomere machinery (Fig. 6). Tay1 homologs appear to be confined to fungi, and database analysis suggests that they emerged prior to the divergence of basidiomycetes from ascomycetes, since family members can be identified in both phyla (e.g., homologs in *Aspergillus* (Pezizomycotina), *Schizosaccharomyces* (Taphrinomycotina), *Puccinia* (Pucciniomycotina), and *Yarrowia* (Saccharomycotina)), but not in more basal branches ^10, 11^. Interestingly, this gene was evidently lost in late branching lineages of Saccharomycotina, given the absence of any discernible homologs in e.g., *Saccharomyces* and *Candida* spp. Indeed, not only is the existence of Tay1 homologs in basidiomycetes and ascomycetes variable, but also their participation in telomere regulation. Our results indicate that at least in *U. maydis* (and possibly in other basidiomycetes), this protein promotes telomere replication and regulates telomere recombination, but is not essential for telomere protection or maintenance. In contrast, *S. pombe* Tay1 functions primarily as a transcription factor ^9^, and *Y. lipolytica* Tay1 seems to be essential for telomere maintenance or protection – *tay1Δ* haploid strains could not be generated, and a *Tay1/tay1Δ* diploid strain exhibited dramatic telomere shortening in comparison to the wild type diploid strain ^10^.

The remarkable plasticity of Tay1 function may be rationalized by considering (1) the relatively recent emergence of the Tay1 protein family in fungi and (2) the drastic alterations in telomere repeat sequence in budding and fission yeasts. One can surmise that prior to the emergence of Tay1 homologs, the functions of duplex telomere-binding protein would be exclusively executed by a TRF/TBF1-like protein in the common ancestor of basidiomycetes and ascomycetes. Once Tay1 evolved to bind telomeres, it could have been enlisted to take over an existing function from the ancestral TRF/TBF1, or it might evolve a new function in telomere regulation. The role of *Um*Trf1 in telomere replication is consistent with either scenario. As a non-essential player at telomeres, the ancient Tay1 would have been relatively unconstrained in adopting other functions, as in the case of SpTay1, which regulates transcription rather than telomeres. On the other hand, the different fate of Tay1 in Saccharomycotina is more easily rationalized with respect to the telomere sequence divergence. In *Y. lipolytica*, the retention of a telomere repeat that contains the canonical GGGTTA sequence enables Tay1 to be telomere-bound and to function in telomere maintenance/protection – though it is unclear why the TRF/TBF1 protein was lost. In the later branching lineages of Saccharomycotina yeasts (e.g., *Saccharomyces* and *Candida*), the complete absence of the GGGTTA sequence renders telomeres refractory to binding by either Tay1 or TRF/TBF1, thus necessitating the utilization of another Myb-containing protein (Rap1) with a more relaxed sequence specificity to bind and protect telomeres ^12^. While TBF1 in these organisms might have been retained because of its roles in transcription and subtelomere regulation ^30, 31, 32^, Tay1 was probably lost due to the lack of any critical function. Altogether, the emergence of Tay1 in fungi and its subsequent loss in specific lineages provide interesting illustrations of functional acquisition and loss in evolution by a new protein family. The ability of the protein family members to evolve different telomeric and non-telomeric functions may have provided added flexibility to fungi to meet the challenges posed by telomere sequence alterations.

### The roles of TRFs in ALT and ALT-related pathways

Among our findings is the demonstration of a stimulatory function for *Um*Trf1 in the ALT-like pathway in *U. maydis*. This is to our knowledge the first example of a telomere protein exerting a positive regulatory function in ALT-related pathways, and could have relevance for similar pathways in mammalian systems. Both mammalian TRF1 and TRF2 have been shown to bind and regulate the cognate BLM, and BLM is strongly implicated in promoting ALT and telomere BIR (break-induced replication)^5, 33, 34, 35^. While these observations hint at a positive regulatory role for mammalian TRFs in ALT, no direct evidence has been reported. Experimental interrogation of this idea is likely to be challenging: To wit, given the roles of TRF1 and TRF2 in telomere integrity and telomere protection, studies that address the interplay between TRF1/TRF2 and BLM in ALT will require appropriate separation-of-function mutants. In this regard, the rather limited telomere function of *trf1* in *U. maydis* allowed us to address the issue more easily through the use of *trf1Δ* null mutations.

The current work also points to potential complications in studies that investigated ALT mechanisms through the use of TRF1-FokI, a fusion protein that induces telomere-specific breaks ^36, 37^. The DNA synthesis/repair triggered by the induction of TRF1-FokI has been proposed to mimic the natural ALT pathway. However, if TRF1-FokI retains the ability to recruit or activate BLM, the breaks generated by this fusion protein might be preferentially processed by BLM, and preferentially channeled toward specific repair pathways rather than those utilized by endogenous ALT substrates. Given the evidence for multiple ALT pathways with distinct factor requirements ^38, 39, 40^, the roles of telomere proteins in regulating these pathways and in modulating the outcomes of BIR-like DNA synthesis at telomeres would appear to warrant further investigation.

## Methods

### *Ustilago maydis* strains and growth conditions

Standard protocols were employed for the genetic manipulation of *U. maydis* ^41, 42, 43^. All *U. maydis* strains used in this study were haploid and were derived from the UCM350 or FB1 background ^42, 44^ and are listed in Supp Table 1.

The *trf1Δ* and *ku70^nar1^ trf1Δ* mutants were constructed by replacing the entire *trf1* open reading frame in FB1 and *ku70^nar1^* with a cassette expressing resistance to hygromycin (Hyg^R^) through homologous recombination as described previously ^17, 42, 45^. Briefly, the plasmid harboring the *trf1* disruption cassette with the *HYG^R^* marker (named pUTrf1-Hyg) was constructed through Gibson Assembly (NEB) of four overlapping PCR fragments, which were generated using appropriate primers (Table S2) and either genomic DNA or plasmid pUMar261 as templates. The four PCR fragments linked by Gibson assembly include: (i) 5’ UTR of *trf1* connected to the 5’ end of the hygromycin ORF, (ii) the complete hygromycin ORF flanked by short segments from the 5’ and 3’UTR of *trf1*, (iii) 3’ end of the hygromycin ORF linked to 3’ UTR of *trf1*, and (iv) the vector sequence from pUMar261 flanked by short segments from the 5’ and 3’UTR of *trf1*. The disruption cassette was generated by cleaving pUTrf1-Hyg with *Eco*RI and *Bam*HI, and used for transformation. The correct mutant strains were identified by PCR analysis.

The *trf2^crg1^* strain was constructed by integrating linearized pRU-trf2 plasmid containing fragments that match (1) 700 bp of the *trf2* promoter and (2) the first 700 bp of the *trf2* ORF into the *trf2* genomic locus in UCM350. Briefly, the two *trf2* fragments were generated by PCR using appropriate oligos with flanking restriction enzyme sites (Table S2), and then cloned together in between the *Nde*I and *Eco*RI sites of pRU12 ^24^. Correct recombination of *Xba*I treated pRU-trf2 into the *trf2* locus will lead to the insertion of the *CBX^R^* marker and the *crg1* promoter precisely adjacent to and upstream of the *trf2* ATG initiation codon. Following transformation, the correct mutant strains were selected on YPA supplemented with carboxin (4 μg/ml) and confirmed by PCR. The strains were grown in YPA (1% yeast extract, 2% peptone, 1.5% arabinose) or YPD (1% yeast extract, 2% peptone, 2% dextrose) to induce or repress *trf2* expression, respectively.

### Telomere length and structural analyses

Southern analysis of telomere restriction fragments (TRF) was performed using DNA treated with *Pst*I or *Alu*I plus *Rsa*I as described previously ^17, 19^. The blots were hybridized to a cloned restriction fragment containing 82 copies of the TTAGGG repeats. The in-gel hybridization assays for detecting ssDNA on the G- and C-strand have also been described ^16^. STELA for telomeres bearing UT4 subtelomeric elements was performed essentially as reported before ^21^. To detect telomere-telomere fusions, chromosomal DNA was subjected to PCR using two subtelomeric primers (UT4-2116-F and UT6-2210-F) that are extended by DNA polymerase towards the chromosome ends (Fig. 6C). The PCR reactions (20 μl) contained 1x Failsafe PreMix H, 0.1 μM each of UT4-2116-F and UT6-2210-F, and 2 U Failsafe™ polymerase, and the thermocycling consisted of 36 cycles of 94°C for 30 sec, 63°C for 30 sec, and 72°C for 2 min. The fusion PCR products were detected by Southern using an UT6 subtelomeric probe ^21^. C-circle assays were carried out by the rolling circle amplification method ^22^, with modifications that were also previously described ^19^.

### Purification of Trf1-FG, Trf2ΔN-FG3, and Blm-FG

To express full length *U. maydis* Trf1 and Blm, we generated FLAG-tagged open reading frames from genomic DNA by PCR amplification using suitable primer pairs (Table S2). The PCR fragments were cloned into the pSMT3 vector to enable the expression of His_6_-SUMO-Trf1-FG and His_6_-SUMO-Blm-FG fusion proteins. For the N-terminally truncated Trf2 (amino acid 415 to 1528), the PCR fragment was inserted into a modified pSMT3 vector carrying a FG_3_ tag downstream of the cloning site. BL21 codon plus strains bearing the expressing plasmids were grown in LB supplemented with kanamycin and induced by IPTG as previously described ^46^. Each fusion protein was purified by (i) Ni-NTA chromatography, (ii) ULP1 cleavage and, and (iii) anti-FLAG affinity chromatography as previously described ^47^.

### EMSA Assays

For DNA binding assays, purified Trf1-FG or Trf2ΔN-FG_3_ was incubated with 1-10 nM P^32^-labeled probe and 0.5 μg/μl BSA in binding buffer (25 mM HEPES-KOH, pH 7.5, 5 % glycerol, 3 mM MgCl_2_, 0.1 mM EDTA, 1 mM DTT) at 22^0^C for 15 min, and then subjected to electrophoresis through a 6 % non-denaturing polyacrylamide gel in 0.2x TBE followed by PhosphorImager analysis.

### Helicase Assays

The assays (modified from ^33^) were performed in 20 μl of helicase buffer (20 mM Tris.HCl, pH 7.5, 2 mM MgCl_2_, 0.1 mg/ml BSA, 1 mM DTT, 2 mM ATP) containing 0.4 nM labeled dsDNA and indicated amounts of Blm-FG, Trf1-FG, and Trf1^N274^-FG. After 20 min of incubation at 35^0^C, the reactions were terminated by the addition of 4 μl stop solution (30 % glycerol, 50 mM EDTA, 0.9 % SDS, 0.1 % BPB and 0.1 % xylene cyanol) and 20 ng of unlabeled bottom strand oligo (as trap to prevent reannealing of unwound ssDNA). The products were separated by 10% polyacrylamide gel (acrylamide : bis-acrylamide = 29 : 1) containing 1x TBE, and then quantified by PhosphorImager analysis.

## Acknowledgments

We thank Qingwen Zhou for purified Rad51 and Brh2/Dss1, and members of our laboratories for discussion. This work was supported by a pilot project grant from the Sandra and Edward Meyer Cancer Center (N.F.L. and W.K.H.) and NSF MCB-1817331 (N.F.L.).

## Author Contributions

EYY, WKH, and NFL conceived the study. EYY, SZ and NFL designed and executed the experiments with contributions from MH and JS. EYY and NFL interpreted the data and wrote the manuscript with advice from SZ and WKH.

