## Supplemental Figures and Tables for "Structurally distinct duplex telomere repeat-binding proteins in *Ustilago maydis* execute specialized, non-overlapping functions in telomere recombination and telomere protection"

Yu et al.

Supp. Fig. 1

A

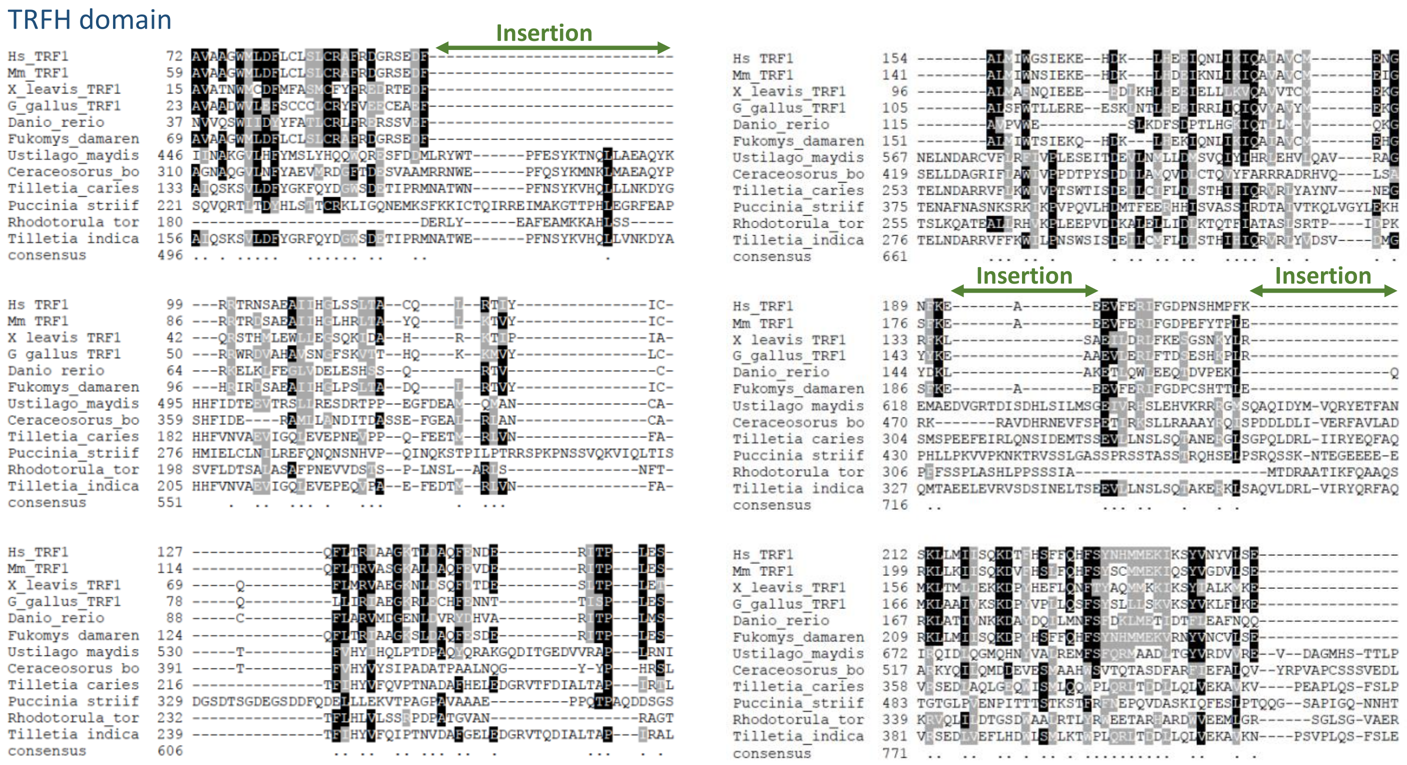
TRFH domain
Insertion
Insertion
Insertion

B

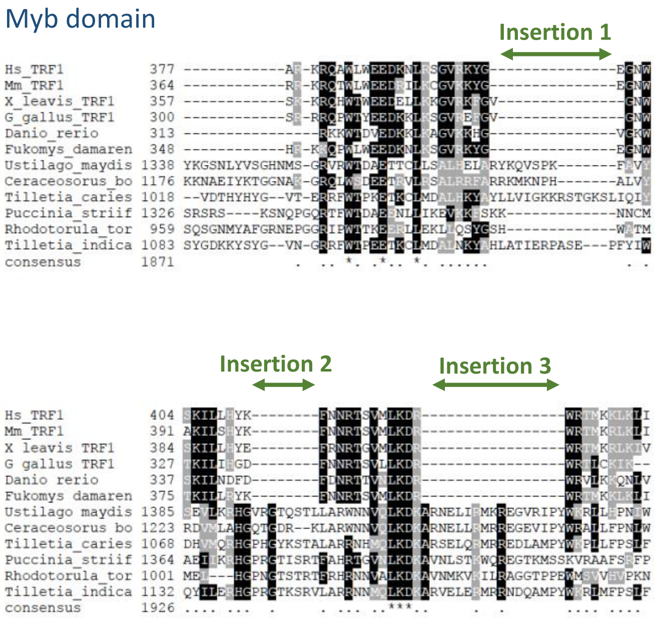
Myb domain
Insertion 1
Insertion 2
Insertion 3

**Supp. Figure 1. Multiple sequence alignments of the TRFH and Myb domain in metazoan TRF1 and Fungal TRF/TBF1-like proteins**

(A) Full length proteins were aligned using T-coffee and the alignment for TRFH region is displayed. The homologs include: Hs_TRF1, from *H. sapiens*; Mm_TRF1, from *M. musculus*; X_leavis_TRF1, from *X. leavis*; G_gallus_TRF1; from *G. gallus*; Danio_rerio, from *D. rerio*; Fukomys_damaren; from *F. damarensis*; Ustilago_maydis; from *U. maydis* (UMAG_02458); Ceraceosorus_bo, from *C. bombacis* (CEH14748); Tilletia_caries, from *T. caries* (OAJ23777); Puccinia_striif, from *P. striiformis* (KNF05609); Rhodotorula_tor, from *toruloides* (XP_016271506); Tilletia_indica, from *T. indica* (OAJ06516). Three regions of insertions in the fungal proteins are highlighted.

(B) Same as in A except that the Myb domain alignment is displayed. The three regions of insertions in the fungal proteins are highlighted.

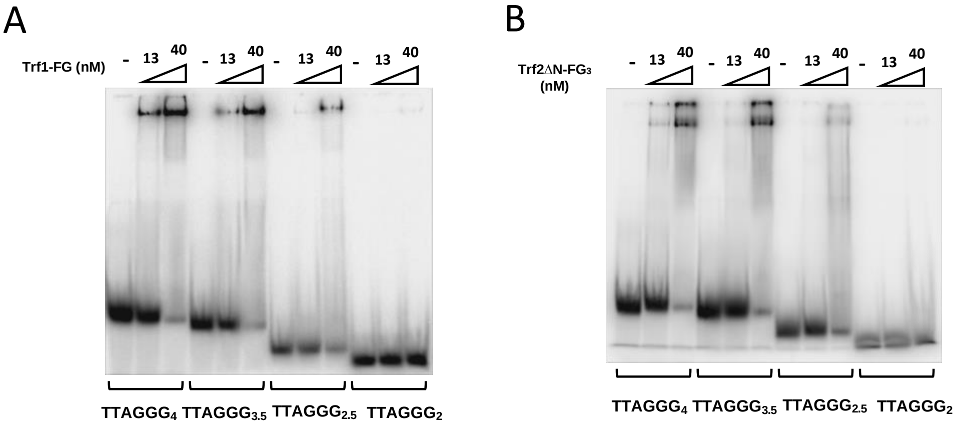

| | |
|---|---|
| **TTAGGG$_4$** | TTA GGGTAA GGGTAA GGGTAA GGG |
| **TTAGGG$_{3.5}$** | TTA GGGTAA GGGTAA GGGTTA G |
| **TTAGGG$_{2.5}$** | TTA GGGTAA GGGTAA G |
| **TTAGGG$_2$** | GGTAA GGGTAA G |

**Supp. Figure 2. Analysis of the minimal DNA target site for *Um*Trf1 and *Um*Trf2**

(A) EMSA was performed using 5 nM of the telomere probes and the indicated concentrations (in nM) of Trf1-FG.

(B) Same as in A except that Trf2ΔN-FG$_3$ was used instead of Trf1-FG. The sequences of the DNA probes used for these assays are displayed at the bottom.

Supp. Fig. 3

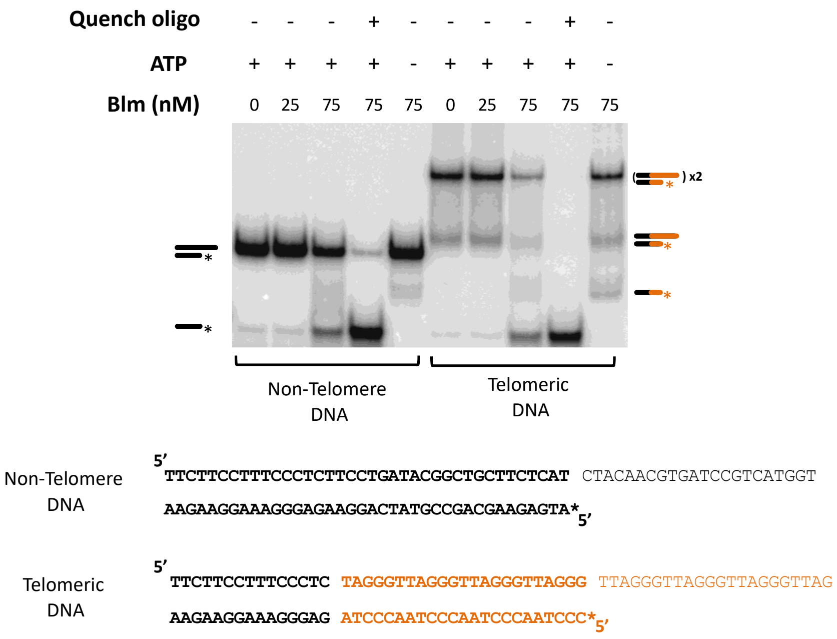

**Supp. Figure 3. Unwinding activity of *U. maydis* Blm helicase on telomeric and non-telomeric DNA**

Helicase assays were performed using the indicated substrates and varying concentrations of Blm in the absence or presence of ATP. The reaction mixtures were analyzed by native gel electrophoresis and PhosphorImager scanning. Labeled species that correspond to dsDNA substrates and ssDNA products are designated schematically to the left and right of the gel. Asterisks indicate the positions of the $P^{32}$ label. The sequences of the substrates are displayed at the bottom, with the non-telomeric and telomeric regions shown in black and brown, respectively.

Supp. Fig. 4

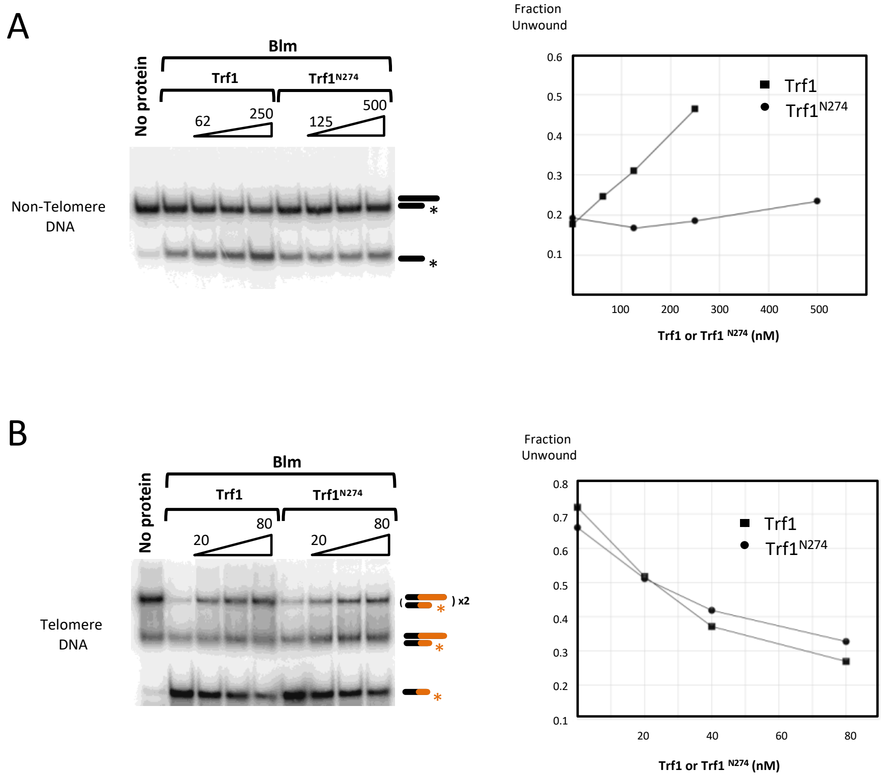

**Supp. Figure 4. The effects of Trf1 and Trf1$^{N274}$ on Blm helicase activity.**

(A) The helicase activity of Blm on non-telomeric DNA was examined in the presence of increasing concentrations of either Trf1 or Trf1$^{N274}$. The assays results are shown on the left and the quantitation plotted on the right.

(B) Same as in A except that assays were performed using the telomeric substrate.

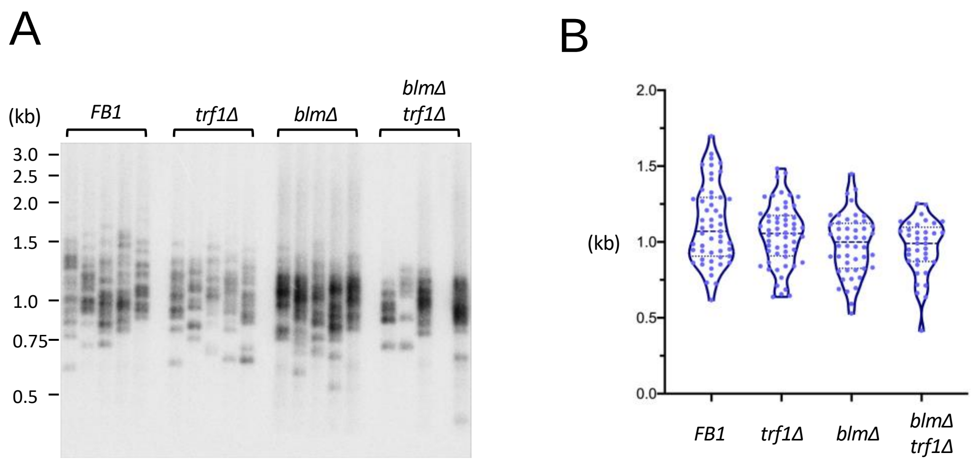

**Supp. Figure 5. STELA analysis of *U. maydis* telomere replication mutants**

(A) STELA assays for the indicated strains were performed in quintuplet. The amplified telomere fragments were analyzed by Southern using a probe that spans a region of the UT4/UT5 subtelomeric element.

(B) The sizes of the STELA fragments in A were analyzed using the TeSLA program [47] and the results shown as violin plots.

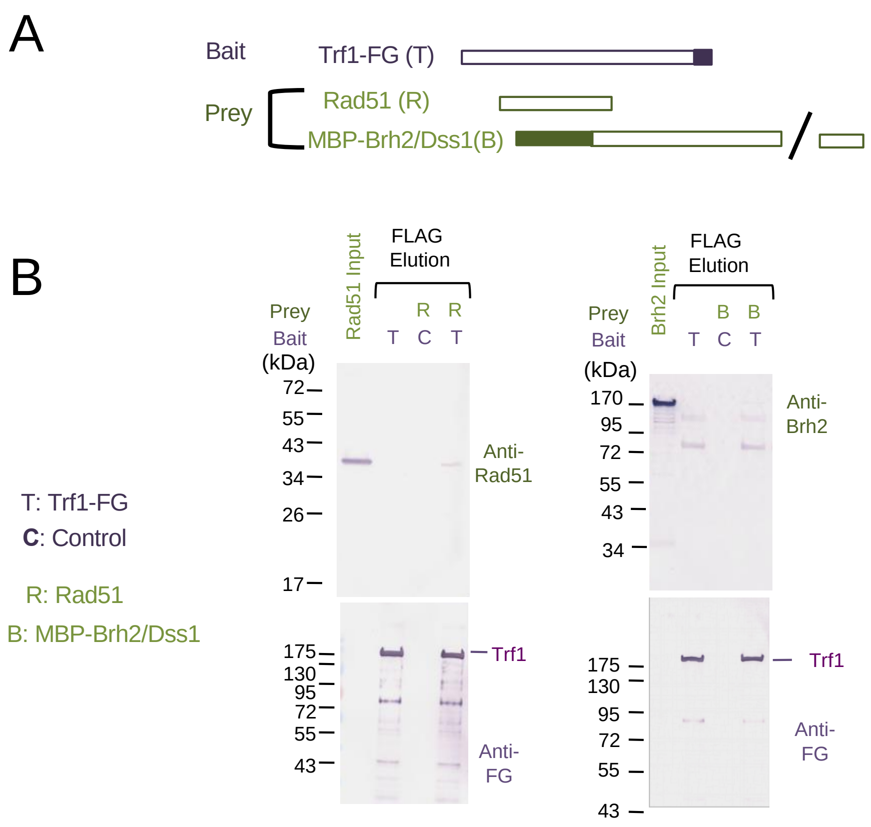

**Supp. Figure 6. Analysis of potential interaction between Trf1 and either Rad51 or Brh2/Dss1**

(A) The Bait and prey used in the pull down assays are schematically illustrated. Fill boxes indicate the locations of the affinity/epitope tags.

(B) Trf1-FG was immobilized on FLAG beads and incubated with either Rad51 or the MBP-Brh2/Dss1 complex (~1 µM). Following incubation, the beads were washed and then eluted with FLAG peptide. The levels of the bait and prey proteins in the elution samples were analyzed by Western using appropriate antibodies. Neither Rad51 nor Brh2 could be detected in the pull down samples, suggesting that they do not interact strongly with Trf1.

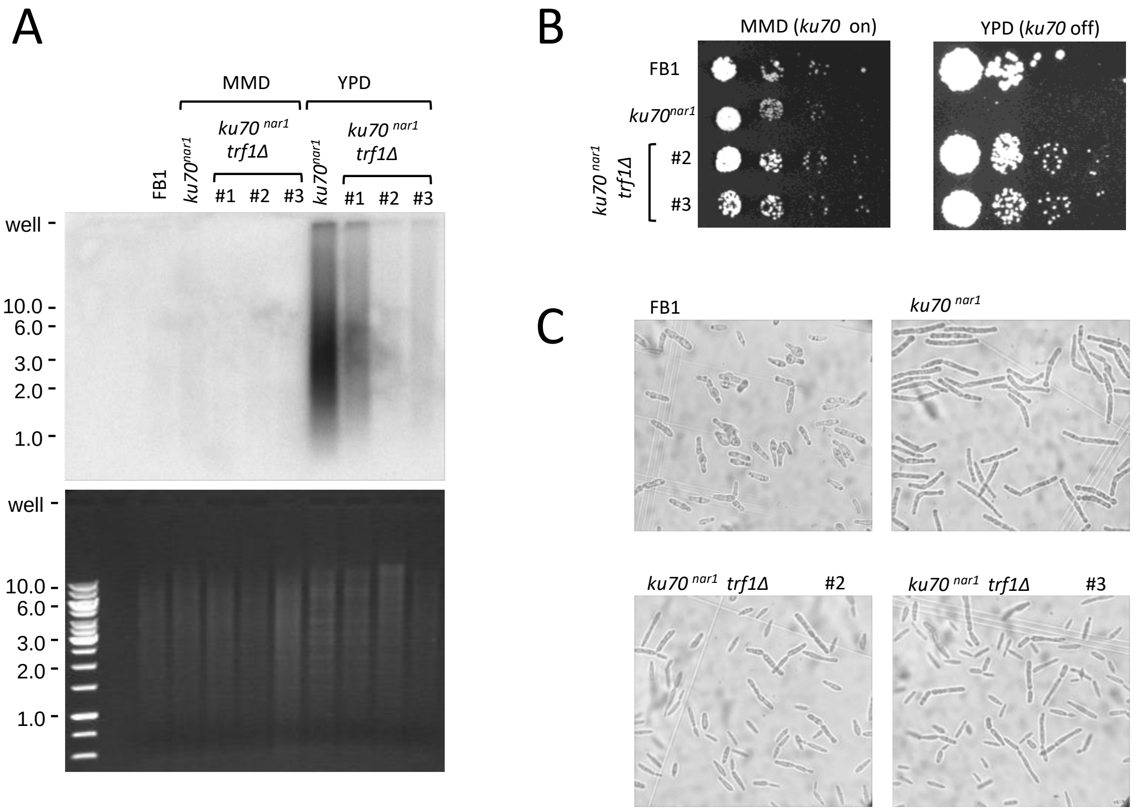

**Supp. Figure 7. The effects of *trf1* deletion on C-strand ssDNA, as well as the growth and morphology of *ku70$^{nar1}$* mutant.**

(A) The levels of C-strand ssDNA in the indicated strains grown in MMD and YPD were analyzed by in-gel hybridization (top). The ethidium bromide stained gel shows that similar amounts of DNAs were analyzed (bottom).

(B) Two independently constructed *ku70$^{nar1}$ trf1Δ* clones were analyzed for growth on MMD and YPD along with the parental FB1 strain and the *ku70$^{nar1}$* single mutant. Serial 6-fold dilutions of cultures of the same density were spotted on the plates, incubated at 30$^{o}$C for 2 ½ days, and then photographed.

(C) The indicated strains were first grown in liquid MMD culture, and then washed and resuspend in YPD such that $OD_{600}$ = 0.05. The cultures were grown for 20 hours at 30$^{0}$C in YPD, and then examined under the microscope.

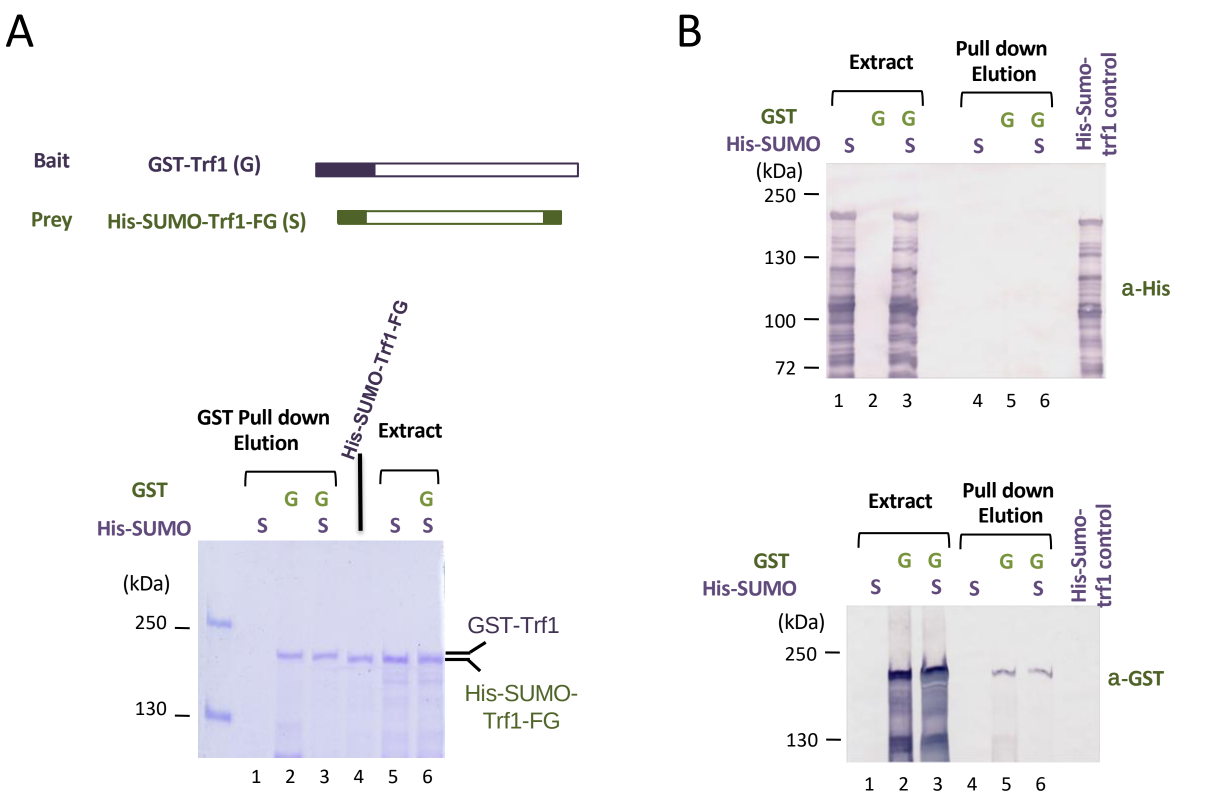

**Supp. Figure 8. Analysis of potential interaction between GST-Trf1 and His-SUMO-Trf1-FG**

(A) Differently tagged Trf1 proteins (GST-Trf1 and His-SUMO-Trf1-FG) were expressed separately or together in *E. coli* and subjected to glutathione-Sepharose pull down. The tagged proteins are illustrated schematically on top. The extracts and pull down samples were analyzed by SDS-PAGE and Coomassie staining, and the stained gel shown on the bottom. Note that the faster migrating His-SUMO-Trf1-FG was expressed at a much higher level than the slower migrating GST-Trf1, but could not be detected in the pull down samples (compare lanes 3 and 6), suggesting that the two proteins did not form a stable complex.

(B) The indicated extracts and pull down samples were subjected to Western analysis using anti-HIS and anti-GST to detected His-SUMO-Trf1-FG and GST-Trf1, respectively. Note that even though there was abundant His-SUMO-Trf1-FG (both full length and proteolytic fragments) in the extracts, none could be detected in the pull down samples (compare lanes 3 and 6), again indicating that the two tagged Trf1s do not form a stable complex.

Supp. Table S1. *U. maydis* strains used in this study

| **Alias (Haploids)** | ***Relevant Genotype*** | **Reference** |
|---|---|---|
| FB1[a] | Wild type | (Banuett and Herskowitz, 1989) |
| UEY25 [ab] | *trf1Δ* | This work |
| UNFL100 [ac] | *blmΔ* | This work |
| UNFL101 [abc] | *trf1Δ blmΔ* | This work |
| UCS33 | $uku70^{nar1}$ | (de Sena-Tomas et al., 2015) |
| UEY26 [ab] | $uku70^{nar1}$ *trf1Δ* | This work |
| UCM350 [d] | wild type | Kojic et al., 2002 |
| UEY15 | *trt1Δ rad51Δ* | (Yu et al., 2018) |
| USZ100 [de] | $trf2^{crg1}$ | This work |

[a] The genotype of FB1 is *a1b1* for the mating type loci.
[b] *trf1* was disrupted by the insertion of *hph* cassette expressing the hygromycin resistance gene ($Hyg^R$).
[c] *blm* was disrupted by the insertion of *cbx* cassette expressing the carboxin resistance gene ($Cbx^R$).
[d] The genotype of UCM350 *is nar1-6 pan1-1 a1b1*. *nar*, *pan*, and *ab* indicate inability to reduce nitrate, auxotrophic requirement for pantothenate, and mating type loci, respectively.
[e] *trf2* was placed downstream of the arabinose-dependent *crg1* promoter through the introduction of a $Cbx^R$-containing cassette.

Supp. Table S2. Oligos used in this study

| Name | Sequence 5' to 3' |
|---|---|
| **Protein Expression** | |
| UmTrf1-F-Hind | AAT AAGCTT CT ATG CCG TCG CAT CCG CAA CCG GCT |
| UmTrf1-R-FG-NotI | AAT GCGGCCGC CTA CTT GTC ATC GTC ATC CTT GTA ATC ATG CTG CTG CCG CTG CTG ATC |
| UmTrf1-274-R-FG-NotI | AAT GCGGCCGC CTA CTT GTC ATC GTC ATC CTT GTA ATC GCG ATC CTT GCG TGC CTT GGA |
| UmBlm-F-Nco | AAT CCATGG CA ATG CCG CAA TCC GCA CTA ACC CCA |
| UmBlm-R-FG-NotI | AAT GCGGCCGC CTA CTT GTC ATC GTC ATC CTT GTA ATC ACC CGA ACG AGG TAG ATT GGG C |
| UmTrf2-F-SalI | ATTAC GTCGAC GT ATG TCA GCT TCT GCT CGG AGC |
| UmTrf2-416F-SalI | ATTAC GTCGAC CG CAA TCC GAA CAG CGG TTA |
| UmTrf2-dn-FG-NotI | AA GCGGCCGC TTA CTT GTC ATC GTC ATC CTT GTA ATC TTC GTT TGA AAG AGA ACT CGA CG |
| **Strain Construction** | |
| UmTrf2(1)Nde | GAAATC CAT ATG TCAGCTTCTGCTCGGAG |
| UmTrf2(700)Xba | ATT TCTAGA CATCCAGAGCCTCCTGGGA |
| UmTrf2(-700)Xba | ATA TCTAGA GCAGTCATCATTATTGCCAT |
| UmTrf2(-1)Eco | ATT GAATTC TGTGGCCCACACGTTACT |
| Trf1-5UTR-Vector-F | CCG GGC CCC CCC TCG AGA ATT CGT CAG TGA AAG AAA AGT GTG AAG CAG TAA GTA TTC |
| Trf1-5UTR-Hyg-R | TGT CAC GCC ATG GT ACT GGA GGC GAT CGT GAG AGG CAA GTG GCA GAC GAG |
| Trf1-3UTR-Hyg-F | GCG GCC GCA TTA ATA CAG GCG CCT TCC GTC GAG TCT TAC TAC CAA TCA CG |
| Trf1-3UTR-Vector-R | GGG CGA ATT GGA GCT CGG ATC CAC CAA TCA AGC CGC TTG TGC CAC GCA CAA CAT GCC |
| Hyg-Trf1-5UTR-F2 | GAT CGC CTC CAG TAC CAT GGC GTG ACA ATT GCG GCC GCA CTC GAG TG |
| Hyg-Trf1-3UTR-R2 | GAA GGC GCC TGT ATT AAT GCG GCC GCA CAG CTT CGC GGC GCA GCA G |
| Vector-Trf1-3UTR-F | CAA GCG GCT TGA TTG GTG GATC CGA GCT CCA ATT CGC CCT ATA GTG AGT CGT ATT AC |
| Vector-Trf1-5UTR-R | CTT TTC TTT CAC TGA CGA ATT CTC GAG GGG GGG CCC GGT ACC AGC TTT TGT TCC C |
| **Helicase Assays** | |
| NT-substrate-top | TTCTTCCTTTCCCTCTTCCTGATACGGCTGCTTCTCATCTACAACGTGATCCGTCATGG T |
| NT-substrate-bottom | ATGAGAAGCAGCCGTATCAGGAAGAGGGAAAGGAAGAA |
| Telo-substrate-top | TTCTTCCTTTCCCTC**TAGGGTTAGGGTTAGGGTTAGGGTTAGGGTTAGGGTTAGGGT TAG** |
| Telo-substrate-bottom | **CCCTAACCCTAACCCTAACCCTA**GAGGGAAAGGAAGAA |
| **PCR, hybridization, and EMSA assays** | |
| $TTAGGG_4$ | TTAGGG TTAGGG TTAGGG TTAGGG |
| $CCCTAA_4$ | CCCTAA CCCTAA CCCTAA CCCTAA |
| $TTAGGG_{3.5}$ | TTAGGG TTAGGG TTAGGG TTAG |
| $CCCTAA_{3.5}$ | CTAA CCCTAA CCCTAA CCCTAA |
| $TTAGGG_{2.5}$ | TTAGGG TTAGGG TTAG |
| $CCCTAA_{2.5}$ | CTAA CCCTAA CCCTAA |

| | |
|---|---|
| TTAGGG$_2$ | GG TTAGGG TTAG |
| CCCTAA$_2$ | CTAA CCCTAA CC |
| Yl TEL G-Strand | TTAGTCAGGG TTAGTCAGGG TTAGT |
| Yl TEL C-Strand | ACTAA CCCTGACTAA CCCTGACTAA |
| Cg TEL G-Strand | TGGGGTCTGGGTGCTG |
| Cg TEL C-strand | CAGCACCCAGACCCCA |
| UT4-2116-F | TCGGGCAACGTTCCATGTCG |
| UT6-2210-F | CTACTACACATCGGTTCAGGC |
